# Two Spore Types in a Marine Parasite of Dinoflagellates

**DOI:** 10.1101/2025.04.10.648098

**Authors:** Jeremy Szymczak, Silvain Pinaud, Irene Romero Rodriguez, Marie Walde, Ehsan Kayal, Catharina Alves-de-Souza, Estelle Bigeard, Benoit Gallet, Martin Gachenot, Sophie Le Panse, Cécile Jauzein, Mickael Le Gac, Johan Decelle, Georg Pohnert, Marine Vallet, Arthur M. Talman, Laure Guillou

## Abstract

Marine alveolates (MALVs) are diverse, primarily parasitic micro-eukaryotes that significantly impact marine ecosystems. The life cycles of most MALVs remain elusive and the role of sexual reproduction in these organisms is a key question that may determine their ecological success. In this study we focus on a widespread dinoflagellate parasite of bloom-forming dinoflagellates, *Amoebophrya*.

After infection, we identified two distinct spores, differing in size, ultrastructure, swimming behavior, lifespan, gene expression, and metabolite composition. The smaller spores serve as infectious propagules, equipped with an apical complex for host invasion. They exhibit a distinct, shorter, and straighter swimming pattern, likely optimized for an extended lifespan while enhancing dispersion and chance for host encounters. Transcriptomic analysis reveals that these smaller spores are primed for efficient protein synthesis upon initiating a new infection.

Conversely, the larger spores cannot infect new hosts and are characterized by the expression of meiotic genes, underscoring their sexual nature. They have a shorter lifespan, exhibit more tortuous movement, along display condensed chromosomes, signaling readiness for mating. Interestingly, infected hosts already express meiotic genes, and a single infected host only produces progeny of the same spore type, suggesting that cell fate is determined prior to spore release.

Our study provides one of the first formal demonstrations of a sexually specialized cell in MALVs. Isolating compatible strains for cross-breeding and understanding how environmental conditions favor each reproductive route are the next key questions for elucidating the ecological success of MALVs in marine waters.

**Significance Statement:** Marine alveolates (MALVs) are ecologically significant parasites that impact carbon cycling, causing major disease outbreaks affecting fisheries and aquaculture, and influencing the dynamics of harmful algal blooms. Despite their diversity and wide host range, much of our knowledge comes from environmental DNA, leaving important aspects of their biology, such as their life cycles, largely unknown. This study provides the first evidence of sexual reproduction in MALVs, linking spore polymorphism to infective or sexual routes. This discovery is crucial as sexual reproduction increases genetic diversity and adaptability, aiding MALVs’ resilience in changing environments. Understanding MALVs’ reproductive strategies deepens our insight into their ecological roles and their broader impact on marine ecosystems.

## Introduction

Sexual reproduction, which emerged approximately 2 billion years ago^1^, is a dominant but not universal process among eukaryotes, and remains particularly underexplored in many lineages, especially among unicellular parasites. Parasitism itself is a key ecological process that enhances ecosystem resilience and exerts evolutionary pressure on both hosts and parasites. Although most unicellular parasites primarily reproduce asexually, some can switch to sexual reproduction as a strategy to increase genetic diversity and maintain infectivity^2^. Given the ubiquity of parasites, understanding the factors that drive, sustain, and limit their diversity and survival is crucial in the fields of ecology and evolution.

In this study, we closely examined the life cycle stages of the parasite *Amoebophrya* sp., which belongs to marine alveolates (MALVs)—an unfathomable and diverse polyphyletic group of early-branching dinoflagellates, discovered within the marine plankton^3–5^. MALVs were identified as one of the most hyperdiverse lineages in the metabarcoding dataset collected during the Tara Oceans expedition^6^. Despite their diversity, only a few species across MALV lineages has been formally described, with most being obligate aplastidial parasitoids intracellular biotrophs (i.e., the host is maintained alive during the infection but killed at last). This diverse assemblage encompasses a broad spectrum of specialized parasites, each adapted to infect specific host types. Collectively however, MALVs can infect a wide variety of marine hosts, ranging from unicellular organisms like dinoflagellates, ciliates, and radiolarians to larger multicellular animals such as crustaceans and fish^5^. MALVs exert substantial influence on marine health and industry: some species cause devastating epizootics impacting fisheries and aquaculture^7^, while others drive the collapse of toxic algal blooms, reshaping ^8,9^phytoplankton community dynamics^8^. They contribute actively to carbon cycling by facilitating organic matter transfer within microbial food webs and enhancing carbon export in nutrient-poor, oligotrophic waters^9^. Despite their ecological importance, MALVs’ life strategies and propagation mechanisms remain poorly understood, highlighting the need for further research into their life cycles and reproductive modes.

Within their hosts, MALVs typically progress from an intracellular feeding stage (trophont) to a reproductive stage (sporont), releasing motile spores with dinoflagellate-like flagella into the water. MALVs produce different spore types that vary in size though their functions remain unclear. For instance, *Syndinium* (MALV Group IV) was found to release three distinct spore types from infected copepods^10^. In *Ichthyodinium chabelardi* (MALV Group I), a parasite of fish embryos and larvae, small invasive spores are produced after three divisions, while larger spores, with unknown functions, are formed after two divisions^11^. Similarly, *Euduboscquella cachoni* (MALV Group I) releases small or large spores from tintinnid hosts, with larger spores evolving into cyst-like forms^12^. Cyst formation mirrors the reproductive behavior of many dinoflagellates, which are haploid during their vegetative phase but produce diploid zygotes after sexual fusion, eventually forming resting cysts in harsh environmental conditions^13^.

This study aimed to uncover the developmental origins and functions of distinct spore types in marine alveolates (MALVs), specifically investigating the relationship between spore size polymorphisms and reproductive strategies, including potential sexual reproduction. We focused on *Amoebophrya* sp. within the *A. ceratii* species complex, which infects dinoflagellates. This organism serves as an ideal model among MALVs because both the host and the parasite are unicellular, with a rapid generation time of 3 to 4 days. The parasite exhibits strong green autofluorescence under blue light excitation, allowing for its detection by flow cytometry and microscopy both in intracellular and free-living stages. Using integrated methods such as advanced microscopy, behavioral analysis, metabolite profiling and transcriptomic analyses, we map two different spore cell types in the *Amoebophrya* life cycle, including one participating in sexual reproduction.

## Results

### *Amoebophrya* trophonts generate two distinct spore types

The *Amoebophrya* sp. strain A120, which infects the dinoflagellate *Scrippsiella acuminata*, releases two distinct spore populations, termed P1 and P2. These populations can be differentiated by their flow cytometry signatures (Fig. 1), with P1 exhibiting lower green fluorescence and forward scatter (FSC) compared to P2. In mixed-infected host populations, P2 spores were first released around 36 hours after inoculation, while P1 spores were released 37.5 hours after inoculation (Fig. S1). After isolating single infected host by micromanipulation, we tracked spore release by flow cytometry. In the majority of cases (41 out of 45), infected host cells produced exclusively either P1 spores (from 6 individuals) or P2 spores (from 35 individuals) (Fig. S2). P1-producing hosts had a significantly higher spore yield, releasing an average of 718 ± 111 spores per host, compared to 190 ± 70 spores from P2-producing hosts. Assuming synchronous divisions of the sporont cells (y = 2n), the number of iterative divisions required to reach these counts was estimated around 10 for P1-producing hosts and 8 for P2-producing hosts.

**Figure 1.**
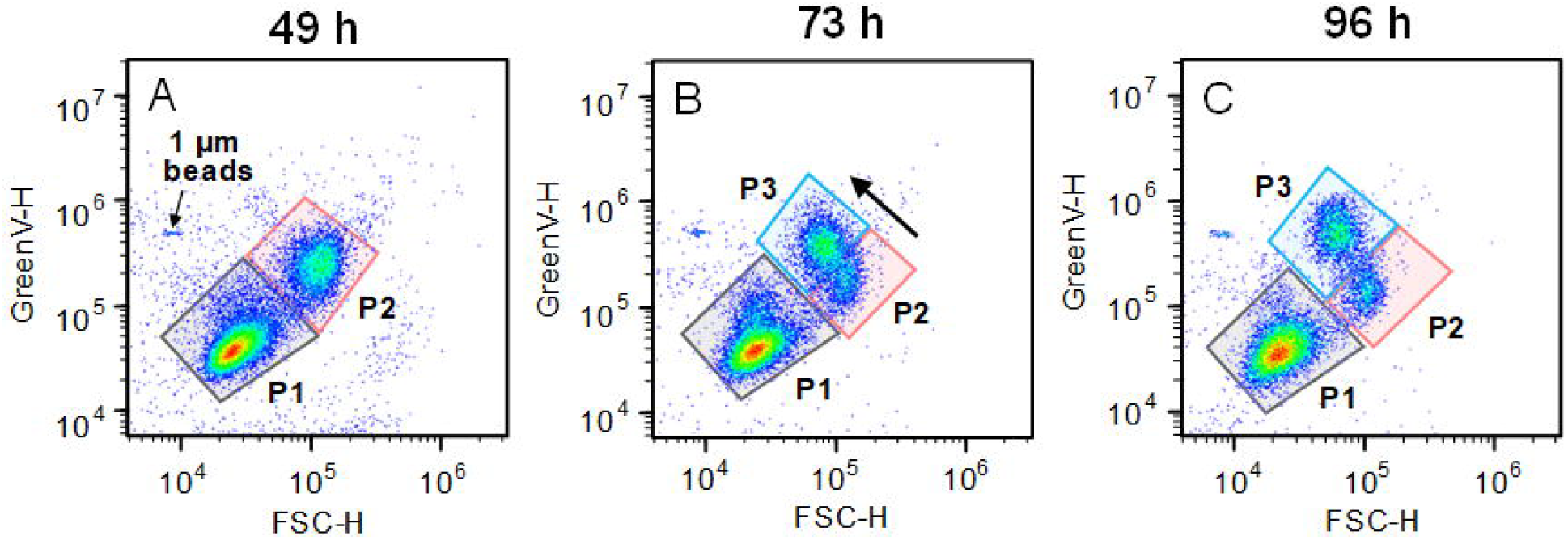
Flow cytometry signatures of the different spore populations—P1, P2, and P3— produced by *Amoebophrya* sp. strain A120 infecting the dinoflagellate *Scrippsiella acuminata*, tracked over time (hours). Differences in green fluorescence (GreenV-H) and forward scatter (FSC-H) for the three populations as observed at (A) 49, (B) 73, and (C) 96 hours post-inoculation. P3 originated from P2 populations 73 hours post inoculation.

The resilience of the two spore populations differed significantly: P1 remained relatively stable in both density and flow cytometry signature for up to 140 hours post-inoculation (the time when the host and parasite were mixed at the start of the experiment). In contrast, the majority, or occasionally the entire P2 population, underwent a notable transformation 50 hours post-inoculation. This transformation was characterized by a reduction in FSC, accompanied by an increase in green fluorescence (Fig. 1). Eventually, a distinct population, P3, emerged (Fig. 1). Both P2 and P3 populations rapidly declined compared to the P1 population (Fig. S3).

### P1 and P2 spores belong to two biologically distinct cell types

Freshly released, P1 and P2 spores can be easily separated based on their size. Using confocal microscopy, we found that P2 spores were nearly twice as large as P1 spores, with an average volume of 27.2 µm^3^ ± 3 µm^3^ (n = 112) compared to 15.5 µm^3^ ± 2.5 µm^3^ for P1 (n = 156) (Fig. S4). P1 spores measured 1.84 ± 0.23 µm in length and 1.37 ± 0.33 µm in width, while P2 spores were larger, measuring 2.9 ± 0.41 µm in length and 2.55 ± 0.25 µm in width, as determined by transmission electron microscopy (Fig. 2). Both spore types possessed two flagella: a larger transverse flagellum with hairs and a smaller hairless longitudinal flagellum (Figs. 2A-B). The lengths of the transverse flagella were similar between the two spore types, measuring 9.48 ± 2.55 µm for P1 and 10.46 ± 2.27 µm for P2. However, the smaller flagellum was generally only visible in P2, measuring 2.11 ± 1.34 µm.

**Figure 2.**
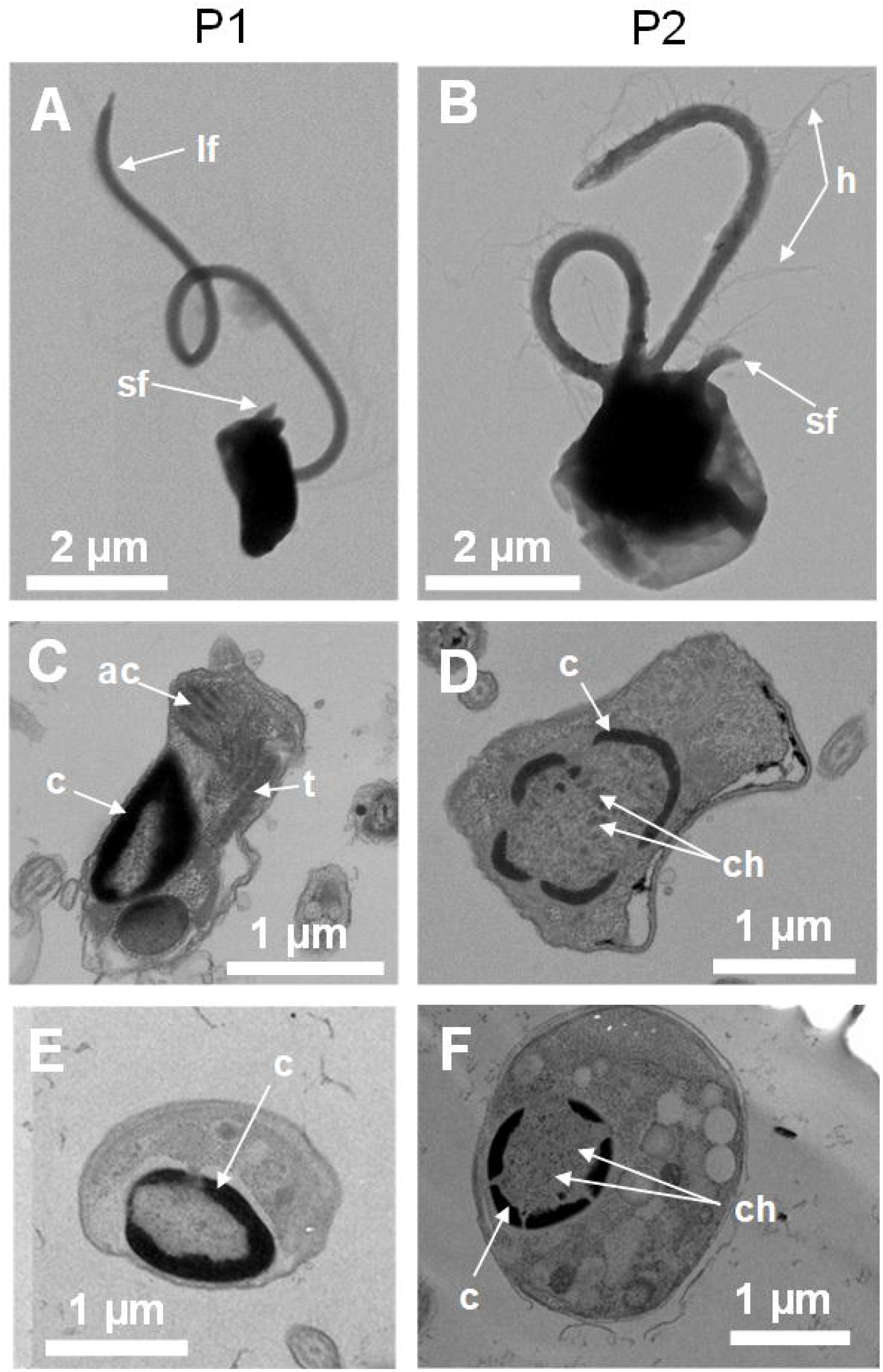
Transmission electron microscopy (TEM) employed to examine P1 (left) and P2 (right) spores using various techniques: (A-B) TEM negative staining, (C-D) thin section with chemical fixative, (E-F) thin section with cryofixation. ac: apical complex with vesicles, c: condensed chromatin, dv: dense vesicles, h: flagellar hairs, ld: lipid droplet, lf: long flagellum, sf: small flagellum, t: trichocyst.

In terms of swimming behavior, P2 spores were significantly more likely to swim (42% of P2 spores versus 24% of P1 spores, p =8.4e-07, odds ratio = 2.3, 95% CI: 1.7-3.4, Fisher-Exact Test). P1 and P2 swam at similar average speed (median of 160 µm/s for P1 and 152 µm/s for P2, p=0.086, Wilcoxon rank sum test) (Fig. 3A). However, P2 spores swam for longer periods (median of 1.26 s for P2 compared to 0.64 s for P1, p=0.004, Wilcoxon rank sum test) and covered greater distances (median of 203 µm for P2 vs. 117 µm for P1, p=0.001, Wilcoxon rank sum test) (Figs. 3B-C). P1 and P2 do not swim in straight lines and P2 swimming tend to be more tortuous than P1 (median tortuosity of 1.9 for P1 and 2.8 for P2, p = 0.005, Wilcoxon rank sum test) (Fig. 3D).

**Figure 3.**
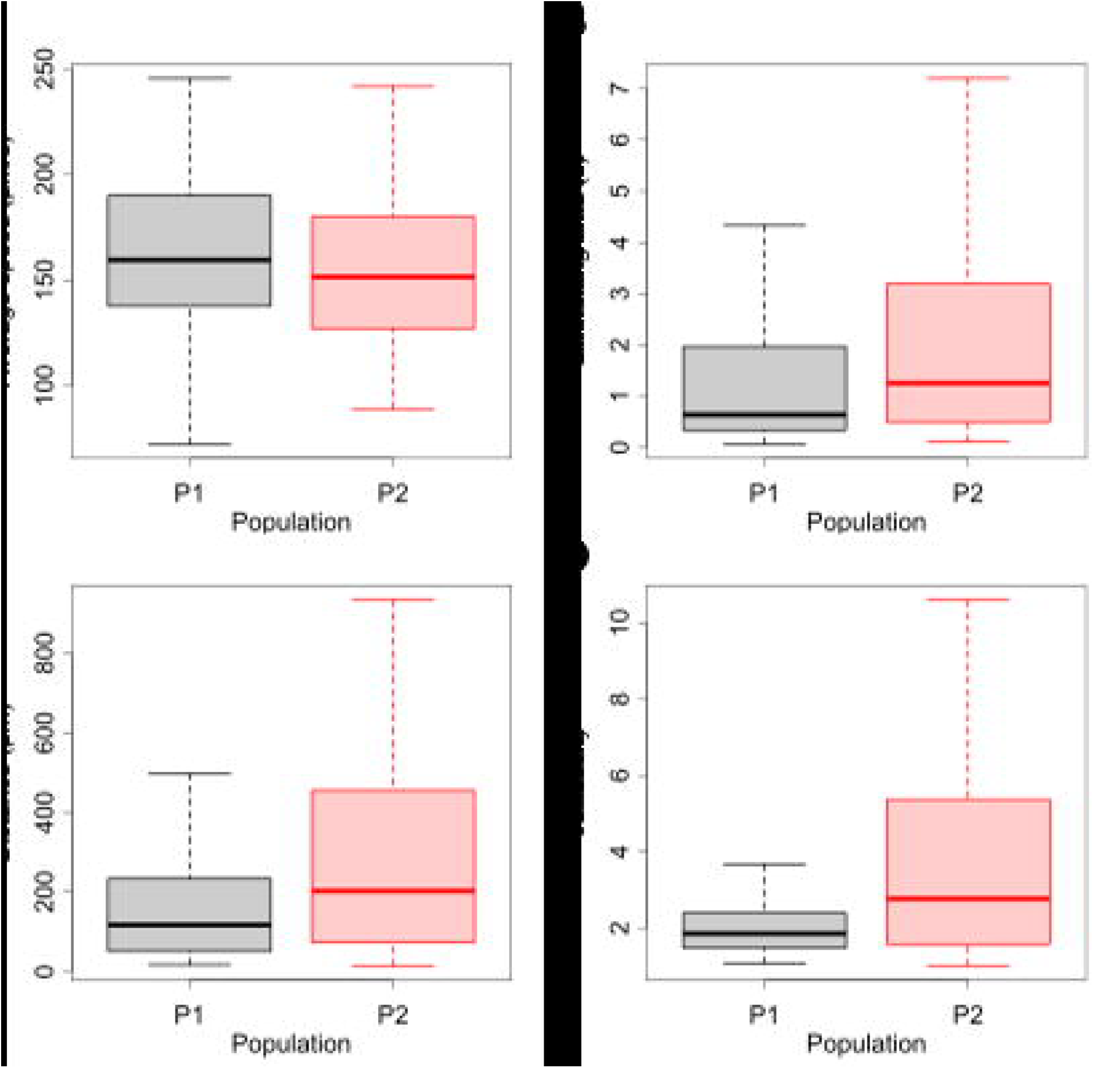
Swimming behavior of P1 and P2 spores. A: average speed, B: Swimming time, C: Distance covered, D: Tortuosity.

Transmission electron microscopy (TEM) revealed differences in nuclear organization: P1 spores exhibited condensed chromatin forming a ring peripherical to the nuclear envelope (Fig. 2C-D), while P2 spores displayed condensed, differentiated chromosomes (Fig. 2E-F). P1 spores had an apical complex-like structure, including vesicles, rhoptry-like organelles, and trichocysts (Fig. 2C-D). In contrast, P2 spores contained numerous droplets in their cytoplasm but no apical complex-like structure, leading to the hypothesis that P2 spores were not infective (Fig. 2F). To test this, we infected cells with a pure culture of P2 spores and monitored this culture over a week. During this time, all P2 spores transitioned to P3, but no infection occurred, leading to the conclusion that neither P2 nor P3 were infective (Fig. S5). We further analyzed the DNA content of mixed spore populations (P1 with P2 and P1 with P3) using flow cytometry (Fig. S6). SYBR green fluorescence intensity ratios were nearly identical for both mixed populations, at 1.17 for P1/P2 and 1.18 for P1/P3, indicating that all three spore types (P1, P2, and P3) share the same ploidy level. A ratio close to 1 for both P1/P2 and P1/P3 nuclei fluorescence indicates a similar level of ploidy for the three spore morphotypes. The results remained unaffected by a heating treatment of 70°C for 20 minutes (not shown).

### One of the spore types is associated with meiosis

To gain deeper insight into the characteristics of the different spore types, we performed transcriptomic analyses on pools of 20 sorted infected host cells at various stages and 10 sorted spores from P1, P2, and P3 populations (Fig. S7). After quality control, we retained transcriptomic data for 10 pools early-infected host cells (i1), 20 pools mid-infected cells (i2/i3), 10 pools late-infected cells (i4), and a total of 41 P1, 15 P2, and 30 P3 spore pools. Mapping reads to a combined host-parasite reference genome-transcriptome, respectively, showed a progressive accumulation of parasite RNA as infection advanced. In late-stage infections, parasite RNA made up ∼75% of total reads (Fig. 4A). Dimensionality reduction based on parasite transcripts revealed distinct expression profiles for different infection stages and spore types, with the exception of mid-infected host cells (i2/i3), which overlapped. P2 and P3 spores were also highly similar and grouped together in subsequent analyses (Fig. 4B). A differential gene expression analysis between P1 and P2+P3 spores identified 2,225 upregulated and 252 downregulated genes in P2+P3 spores.

**Figure 4.**
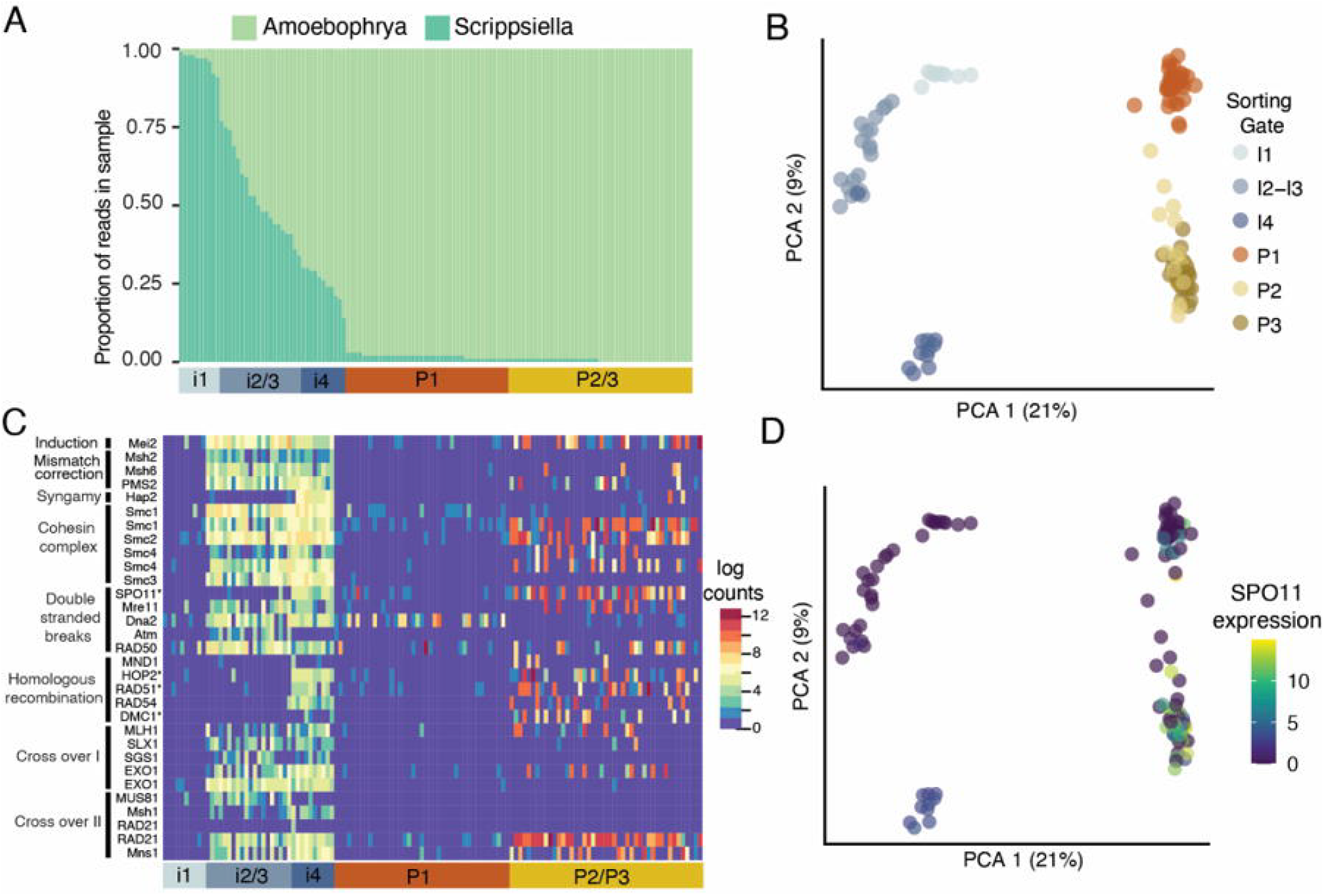
Transcriptomic analysis of parasite populations sorted by flow cytometry. (A) Proportion of reads in sample mapping to either the host *Scrippsiella acuminata* or the parasite *Amoebophrya* sp. strain A120; (B) Principal component analysis (PCA) based on parasite transcriptional profiles of the intracellular stage (trophont) and spores; (C) Heatmap of expression of genes involved in meiosis; (D) An example of gene expression (SPO11, involved in meiosis) over sample (PCA analysis). i1, i2/3, i4: different trophont developmental stages over time. P1, P2, P3: spore populations.

We identified 32 conserved meiotic genes in *Amoebophrya* (Table S1), 21 of which were highly expressed in P2+P3 spores but nearly absent in P1 (Fig. 4C). These included key meiosis-specific genes like SPO11, HOP2, DMC1, and MND1 (Fig. 4D), whose homology has been confirmed in dinoflagellates (Lin et al., 2022). For instance, SPO11, HOP2, and DMC1 are crucial for meiotic processes such as double-strand DNA breaks and chromosomal cohesion, while, the gamete fusogen HAP2 was found only in P2 spores. Meiosis-related genes were already expressed in mid to late intracellular stages (i2/i3-i4), suggesting that sexual reproduction may be already engaged before spore release (Fig. 4C).

In addition to meiotic pathways, other gene sets related to DNA replication were highly upregulated in P2+P3 spores. Genes involved in DNA replication, such as DNA primase, polymerase (alpha and epsilon), topoisomerases I, II, and III, helicases, and DNA replication licensing factors (e.g., MCM proteins), were expressed at significantly higher levels in these spores (Fig. S8). Furthermore, genes related to chromosomal architecture, such as those encoding histones, kinetochore proteins (e.g., NDC80, CDC20, EB1), tubulin, and basal body proteins, were also upregulated, indicating substantial chromosomal reorganization during this stage.

P1 spores appear less transcriptomically active in comparison to P2 and P3 spores, with the notable exception of ribosomal proteins, which were highly expressed in P1 spores, suggesting active ribosome biogenesis. These ribosomal proteins included small nucleolar ribonucleoproteins (snoRNPs) involved in the initial stages of ribosome assembly, along with 12 other ribosomal proteins, many of which were also expressed in P2+P3 spores (Fig. S8).

### Spore type have distinct metabolic profile

We performed a comprehensive metabolite analysis on both exudates (exometabolome, Fig. S9) and cell extracts (endometabolome, Fig. S10) from cultures primarily containing P1, P2, or P3 spores at specific time points (49 hours post infection for P1 and P2, 71 hours post infection for P3). Principal component analysis (PCA) revealed that P2 and P3 spores exhibited similar metabolic profiles, both in their endometabolome and exometabolome, indicating metabolic similarity. Thus, for subsequent analyses, P2 and P3 spores were grouped together. We observed significant differences between the metabolites associated with P1 spores and the combined P2+P3 spores. A curated set of 44 metabolites showed clear distinctions between these spore types. P1 spores displayed a distinct set of metabolites. Five key molecules were identified with high confidence, including isocitric acid, leucine, glycine betaine, indoline, and phenylalanine. Five metabolites were uniquely associated with P2+P3 spores in the endometabolome, with two also found in the exometabolome, suggesting potential extracellular secretion and perhaps cell-cell communication.

## Discussion

In this study, we unveil the distinct functional roles of different spore types produced by the parasite *Amoebophrya sp. strain A120*. Dimorphic spore production is a well-documented feature in most MALVs; however, the specific roles of these spores have remained elusive until now. We observed that a single infected host cell predominantly produces a single spore morphotype, a phenomenon previously documented in other Syndiniales species (e.g., Syndinium^10^, *Ichthyodinium*^11^, *Euduboscquella*^12^). As observed in *Ichthyodinium*^11^, the spore dimorphism found in *Amoebophrya* likely depends on the number of iterative divisions during sporulation, which arises from two distinct pathways of development (8 and 10 rounds of division for P2 and P1, respectively). The two fewer rounds of division could explain the larger cell size of P2 and their earlier production compared to P1. Since P2 is released 1.5 hours before P1, we estimate the duration of each of the two last replications to be about 45 minutes, although replication time may vary in each round. For instance, different replication times were reported for *Duboscquela melo*^14^, but not for *Ichthyodinium chaberladii*^11^.

Our findings indicate that the smaller spore type (P1) serves as the infectious propagule. The presence of an apical complex, as observed in TEM—a characteristic typically found in Apicomplexa parasites —indicates that these spores are equipped to invade the next host. Interestingly, P1 spores swim less often and for shorter period than larger non-infective spore, a strategy that likely minimizes energy expenditure, thereby contributing to their prolonged survival, while their less tortuous swimming trajectories that may be attributed to the reduction of the transversal flagellum.

Transcriptomic analysis revealed that P1 spores had a distinct profile compared to P2+P3 spores. In particular, ribosomal pathways involving small nucleolar ribonucleoproteins (snoRNPs) are pre-activated, preparing P1 spores for rapid protein synthesis following invasion. TEM confirmed highly condensed chromatin peripherical to the nucleus in P1 spores, consistent with lower gene expression. In P1 spores, metabolic profiling identified an accumulation of key molecules from central metabolism, including isocitric acid, leucine, and phenylalanine. As *Amoebophrya* cannot synthesize these compounds *de novo*, they likely acquired from the host. Isocitric acid, a key component of the tricarboxylic acid cycle (TCA) and plays a pivotal role in energy release through the oxidation of acetyl-CoA derived from carbohydrates, fats, and proteins. This finding aligns with the incomplete nature of the TCA in *Amoebophrya*^15,16^, further emphasizing the parasite’s reliance on host-derived metabolites.

P2 gene expression suggests their involvement in the sexual reproduction of the parasite. P2 transform into P3 spores based on their flow cytometric signatures, but this transition does not significantly alter their transcriptomic and metabolomic profiles. Both P2 and P3 spores remain non-infective over time and disappear more quickly compared to the more stable P1 spores. The overexpression of meiotic genes begins during the late stages of intracellular infection, suggesting that the program for sexual reproduction starts well before spore release. This is consistent with the observation that an infected host releases only a single spore type, indicating that the distinction between infection and sexual reproduction is determined early within the host.

P2 spores may act as gametes. Several observations regarding P2 spores align with this hypothesis. For instance, P2 spores swam for longer period than P1 spores, allowing them to cover greater distances while following a more tortuous trajectory, a movement pattern typical for dinoflagellates gametes^17^. Additionally, the chromatin organization within P2 spores, characterized by condensed chromosomes supposedly prepared for pairing, further supports the notion of these spores being primed for sexual reproduction. The identification of specific molecules secreted exclusively by P2 and P3 spores raises the intriguing possibility that they function as pheromones. Such molecules could play a critical role in gamete recognition and fusion.

We observed no zygote formation or changes in ploidy across spore types in our study, suggesting two potential explanations: either meiosis is immediately followed by cell fusion, leading to a transient diploid phase that remains undetectable by flow cytometry, or, more likely, that the *Amoebophrya* sp. strain A120 is heterothallic, relying on cross-breeding between compatible strains for sexual reproduction to succeed. It is worth noting, however, that the activation of the sexual reproduction pathway in heterothallic strains does not necessarily depend on the presence of compatible partners. For instance, in *Alexandrium minutum*, known to be heterothallic, sexual gene upregulation has been documented under nitrogen (N) and phosphorus (P) stress even in monoclonal cultures, suggesting that environmental stressors alone can trigger preparatory pathways for sexual reproduction^18^. This behavior implies that while actual zygote formation requires compatibility, gene expression linked to meiosis and sexual development can be initiated independently of partner availability, possibly as a preparatory response to fluctuating environmental conditions.

In this study, we investigate the functional roles of different spore types in the parasite *Amoebophrya* sp. strain A120, which infects dinoflagellates. We found that smaller P1 spores are infectious, equipped with an apical complex for host invasion, and exhibit distinct metabolic characteristics indicative of host-derived nutrient acquisition. In contrast, larger P2 spores, which arise from fewer rounds of division than P1 spores, are involved in sexual reproduction. The overexpression of meiotic genes in P2 spores and their behavior similar to gametes suggest they play a key role in sexual reproduction. These findings not only enhance our understanding of parasite diversity and survival strategies but also provide insights into the ecological and evolutionary implications of sexual reproduction in parasitic marine systems.

## Materials and Methods

### Culture conditions

The experiments were conducted using cultures of *Amoebophrya* sp. strain A120 (RCC4398), infecting the dinoflagellate host *Scrippsiella acuminata* strain ST147 (RCC1627). Both strains are available at the Roscoff Culture Collection (https://www.roscoff-culture-collection.org/). For most of our experiments, F/2 media was prepared with Red Sea Salt (Red Sea Company) diluted with milli-Q water to achieve a salinity of 27 PSU^19^. The exception to that was for cultures used for metabolomics analyses, for which medium was prepared with natural seawater from the Penzé Estuary collected at a salinity of 27 PSU complemented with 5% soil extract^20,21^.

Stock cultures of both host and parasites were grown at 21°C in 50-mL vented flasks (Culture One) under continuous light conditions at an intensity of 100 µEinstein m2 s−1. Cultures were transferred twice a week, with a volume ratio of 1:4 and 10:1 for the host:medium and parasite:host, respectively.

Except when specified, parasite cultures were synchronized previous to experiments^22^. Briefly, old spores from the cultures were removed which resulted in all newly released spores being of the same age. To do so, infected hosts at the beginning of infection (7-10 hours following parasite inoculation) were gently collected by gravity filtration on 5-10 µm nylon filters (Merck Millipore). Afterward, the filters with the host cells were rinsed on the same support device using a sterile culture medium to ensure the removal of the remaining old spores, and the cells retained on the filter were then collected and diluted into a fresh medium in a new flask.

### Confocal microscopy

Cell volumes in two cultures of freshly released spores were compared, comprising approximately 88% P1 (sample 1) and 85% P2 (sample 2). Confocal microscopy Samples were fixed in a combination of paraformaldehyde (Electron Microscopy Sciences, ref. 15714, 1% final conc.) and EM grade glutaraldehyde (Sigma-Aldrich G5882; Merck, Germany; 0.25% final conc.)^23^ during 15 min at 4°C and mounted into chambered cover glasses (Nunc Lab-Tek II; Merck, Germany). Cells were coated with a fluorescent surface label consisting of 0.1 mg ml−1 poly-L-lysine (PLL; Sigma-Aldrich P5899; Merck, Germany) conjugated with Alexa Fluor 546 (AF546SE, Invitrogen A20002; Thermo Fisher Scientific, MA, USA)^24^. Cells were then imaged directly inside the chambers. 3D confocal images were acquired on a motorized and inverted SP8 laser scanning confocal microscope (Leica Microsystem, Germany) equipped with 63× oil NA 1.4 immersion objective by a semi-automated two-step procedure^25^. After manual detection of cell positions, multichannel fluorescence z-stacks of 155 individual cells in sub-sample 1 and 172 individual cells in sub-sample 2 were automatically recorded (AF546: Ex 552 nm / Em 570-590 nm). 3D volumes were rendered from the AF546 fluorescence signal of the surface cover with Imaris 3D image visualization and analysis software (Oxford Instruments, UK). After the normal distribution of the biovolumes (Shapiro–Wilk test) was validated, the 99% confidence interval was calculated (mean +/-2.576*Standard deviation) to separate spore populations.

### Cell counts and cell sorting by flow cytometry

The infection dynamic was followed using a NovoCyte Advanteon flow cytometer (ACEA Biosciences, San Diego, CA, USA) equipped with blue and violet lasers (488 and 405 nm, respectively)^26^. Sorted populations for microscopy analyses, ploidy level determination, infectivity assessment, and transcriptomic analyses were done using an Aurora CS (Cytek, CA, USA) equipped with three lasers (405, 488, and 640 nm). Culture medium was used as sheath liquid for most experiments, while sterile PBS 1X was employed for transcriptomic analyses. During the sorting process, a threshold of 5,000 FSC was set to ensure accurate separation. A flow rate of 45 µL/min was used throughout all sorting procedures to maintain single-cell purity.

For bacterial counts (considered in metabolomics analyses), 1.5-mL aliquots were fixed with grade II glutaraldehyde (Merck Sigma, 0.25% final concentration) for a minimum of 15 minutes and then stored at –80 °C until analysis. Upon thawing, DNA was stained using SYBR Green-I at a final dilution of 1/50,000, following the protocol outlined by Marie et al.^27^.

### Ploidy level

The ploidy levels of the different spore populations were established following the procedure outlined by Marie et al.^27^. Briefly, nuclei were extracted by mixing 50 μL of freshly produced spore with 450 μL of 0.25X NIB buffer, containing SYBR Green-I at a final concentration of 1/5000.

### Estimation of spore production per host

Individual infected host cells were isolated 24 hours post-inoculation, using an inverted microscope (Carl Zeiss Axio Observer 3, x100 magnification) and bright-field observations. These cells were carefully picked using flame-drawn Pasteur pipettes and transferred to separate wells within a 96-well plate. The plate was kept under the same environmental conditions as the stock cultures and was sampled 1 or 2 days after isolating the host cells. For each well sampled, the total volume of the well (about 100 µL) was analyzed by flow cytometry in order to assess the flow cytometry signature and quantity of produced spores.

### Electron microscopy

For TEM negative staining, 20 µL of freshly released spore were deposited upon a thin carbon film grid and let settle for 20 minutes. Subsequently, grids were incubated for 2 minutes with three droplets of uranyl acetate and then carefully absorbed to remove negative stain excess.

For TEM sections, freshly released spore samples were fixed at 4°C for 24 hours in 25% glutaraldehyde, 0.4 M sodium cacodylate buffer (pH 7.4), and 10% NaCl, then post-fixed at 4°C for 60 minutes in a solution of 1% osmium tetroxide buffered with 0.4 M sodium cacodylate and 10% NaCl. Samples were then dehydrated through an ethanol series (absolute anhydrous ethanol, Carlo Erba) and pellets were embedded in Spurr resin at 60°C for two days. Sections were processed using a diamond knife on a Leica Ultracut UCT ultramicrotome, followed by staining with uranyl acetate and lead citrate. Both negative stained grids and sections were examined and photographed using a JEOL JEM 1400 transmission electron microscope (JEOL, Tokyo, Japan) equipped with a Gatan ultrascan camera.

For TEM after cryofixation, spores were cryo-fixed using high-pressure freezing (HPM100, Leica), followed by freeze-substitution (EM ASF2, Leica) as in Decelle et al.^16^. The freeze substitution mix contained 1% of osmium tetroxide. Ultrathin sections of 60 nm thickness were mounted onto copper grids or slots coated with formvar and carbon. Sections were stained in 1% uranyl acetate (10 min) and lead citrate (5 min). Micrographs were obtained using a Tecnai G2 Spirit BioTwin microscope (FEI) operating at 120 kV with an Orius SC1000 CCD camera (Gatan).

### Swimming behavior

Freshly released spores containing a mix of P1 and P2 populations were used to assess the swimming behavior of spores from five biological replicates. Spore density was adjusted to approximately 300,000 cells mL^-1^. Cells were deposited onto customized chambers made by adding a silicone polymer (Polydimethylsiloxane; PDSM) upon a glass slide (2.5 cm length × 1.3 cm width × 64 µm height = 20.8 µL^3^). They were observed at ×100 magnification using an inverted microscope (Carl Zeiss Axio Observer 3) equipped with a Zeiss Axiocam 705 camera and a LED module for epifluorescence. Spores were observed from their autofluorescence signal, visible using an excitation filter of 420 ± 20 nm and a >470 nm long path emission filter. Videos were recorded with a resolution of 2,464 × 2,056 pixels and a pixel size of 0.548 µm using the ZeissZen blue 3.6 software. Five movies were recorded for each replicate. Movements were tracked for 20 seconds, with a 35 ms exposure time per frame. Cell tracking was performed using the trackpy package (version 0.5.0+3.g3b280ea) in Python, with the following parameters: a minimum size of 23 pixels for a single particle, a maximum displacement of 90 pixels between frames per particle, and a memory of 300 frames to track disappearing particles while maintaining their identification. The minimum mass, representing the expected brightness for a particle, was manually selected for each video based on the level of noise present. The R Statistical Software (version 4.1.2; R Core Team 2021) was used to calculate the following parameters: the percentage of particles in motion (particles moving at least 3 µm between two frames and at least 20 µm over the entire video), as well as, for each particle, the total swimming distance, swimming time, average speed, and tortuosity. This later parameter, calculated by dividing the swimming distance by the shortest distance between the start and end position, indicated how much the swimming deviated from a straight line. Cell sizes were centered by movie. The distribution of these normalized cell sizes, which exhibited a bimodal pattern, was modeled using Gaussian Mixture Modeling implemented in the flexmix R package. Cells were then assigned to P1 or P2 based on their size, using a posterior probability ≥ 0.99. Of the 823 particles tracked, 318 were assigned to P1, and 297 were assigned to P2. The statistical analysis of the generated data was conducted using non-parametric Wilcoxon rank sum tests. A Fisher exact test was used to determine whether the proportion of swimming P1 and P2 differed.

### Test of infectivity

A culture dominated by P2 (100% of total spores) at 773,000 cells mL-1 was filtrated through a 5 µm filter to remove remaining hosts. Then, we initiated infection at different spore: host ratios in single replicate (154: 1, 68: 1, 43: 1, 23: 1, and 14: 1) and using an exponentially growing host (3 days old) with a concentration of 7,800 cells mL–1. Cultures were incubated in 24-well plates, and the parasite prevalence was evaluated by flow cytometry after 24 hours post-inoculation. *Smartseq2 sample preparation, library generation and sequencing*.

For both the spores and the infected host, a total of 10 and 20 cells, respectively, were sorted into 96-well plates (Thermo), each well contained 4 µL of lysis buffer (0.8% of RNAse-free Triton-X (Fisher) in nuclease-free water (Ambion), 2.5 mM dNTPs (Life Technologies), 2.5 µM of oligo(dT) (5’-

AAGCAGTGGTATCAACGCAGAGTACTTTTTTTTTTTTTTTTTTTTTTTTTTTTTT−3’) and 2U of SuperRNAsin (Life Technologies)). Sorted populations are illustrated in the supplementary data 9. Reverse transcription and cDNA amplification were conducted as reported previously^28–30^. Sorted plates were spun at 1,000 g for 10 s and immediately placed on dry ice. Plates were heated at 72°C for 3 min. A reverse transcription mix, containing 1 µM of LNA-oligonucleotide (5’-AGCAGTGGTATCAACGCAGAGTACATrGrG+G-3’; Qiagen), 6 µM MgCl2, 1 M Betaine (VWR), 1X reverse transcription buffer, 50 µM DTT, 0.5 U of SuperRNAsin (Invitrogen), and 0.5 µL of Smartscribe reverse transcriptase (Takara), was added to the plates. The total volume of the reaction was 10 µL. The following cycling conditions were used: a single incubation period at 42°C for 90 min, followed by 10 cycles (42°C/2 min, 50°C/2 min), before a final incubation at 70°C for 15 min. A further PCR mix was added to the plates for whole transcriptome amplification, containing 1X KAPA Hotstart HiFi Readymix (Roche Diagnostics France) and 2.5 µM of the ISO SMART primer^31^ and incubated using the following program: a single incubation at 98°C for 3 min, 30 cycles (98°C/20 s, 67°C/15 s, 72°C/6 min), a final incubation at 72°C for 5 min. Reactions were purified with 1X Agencourt Ampure beads (Beckman Coulter) according to the manufacturer’s instructions. Amplified cDNA was eluted with 10 µL nuclease-free water (Ambion). The quality of a subset of cDNA samples was assessed with the high-sensitivity DNA chip (Agilent) with an Agilent 2100 Bioanalyser. Sequencing libraries were prepared using the Nextera XT 96 kit (Illumina) according to manufacturer recommendations but using quarter reactions. Dual indices set A and B were used (Illumina) for 192 different index combinations, for a total of 192 libraries. Libraries were pooled in two pools and cleaned up with Agencourt Ampure beads (Beckman Coulter) used at a 4:5 ratio. The quality of the libraries was assessed with the high-sensitivity DNA chip (Agilent) ran on an Agilent 2100 Bioanalyser. Both pools were combined and sequenced on a Hiseq 4000 with PE150 (Genwiz, Germany). FASTQ files were obtained after base calling and demultiplexing with Illumina’s software. Nextera adapter sequences were trimmed with cutadapt (v3.4) using cutadapt -a CTGTCTCTTATACACATCT -A AGATGTGTATAAGAGACAG --length 50^32^. A combined reference of A120 and its host *Scrippsiella acuminata*^33,34^ was created using bedtools^35^ and indexed with HISAT2 (v2.2.1)^36^. Reads were aligned to this indexed reference with HISAT2 using hisat2 --max-intronlen 5000 -p 8 -q --very-sensitive. SAM files were converted to BAM using samtools-1.2 view –b and sorted with samtools-1.2 sort^37^. Uniquely mapped reads were selected with Sambamba^38^. BAMs were sorted with samtools sort and reads counted with samtools view. Cells were first filtered based on the number of transcripts detected > 250 and more than > 2,000 reads. After filtration, 126 transcriptomes were used for further analysis. Differential expression analysis was conducted in DESeq2 with default parameters^39^. Differentially expressed gene and log fold changes were visualized with the gplots package (v3.1.3) with heatmap.2.

### Orthologues of sex-related genes

We used the method described in Decelle et al.^16^ to identify orthologues of sex-related genes. Reference proteins of interest listed in previous studies^40–42^ were downloaded from the UniProtKB (https://www.uniprot.org) and VEuPathDB (https://veupathdb.org/veupathdb/app) databases (last access September 2023).

## Acknowledgments

This work used the platforms of the Grenoble Instruct-ERIC center (ISBG; UAR 3518 CNRS-CEA-UGA-EMBL) within the Grenoble Partnership for Structural Biology (PSB), supported by FRISBI (ANR-10-INBS-05-02) and GRAL, financed within the University Grenoble Alpes graduate school (Ecoles Universitaires de Recherche) CBHEUR-GS (ANR-17-EURE-0003). We thank Guy Schoehn and Christine Moriscot, and the electron microscope facility at IBS, which is supported by the Rhône-Alpes Region, the Fondation Recherche Medicale (FRM), the fonds FEDER, the Center National de la Recherche Scientifique (CNRS), the CEA, the University of Grenoble, EMBL, and the GISInfrastructures. For the purpose of Open Access, a CC-BY 4.0 public copyright licence (https://creativecommons.org/licenses/by/4.0/) has been applied by the authors to the present document and will be applied to all subsequent versions up to the Author Accepted Manuscript arising from this submission.

