## Supplementary Figure 1 for "Two Spore Types in a Marine Parasite of Dinoflagellates"

**Supplementary Figure 1:** The complete infection cycle of *Amoebophrya sp. strain A120* within *Scrippsiella acuminata* monitored by flow cytometry. Infected host cells can be distinguished from healthy cells based on their green autofluorescence content, which increased by a factor of 100× during parasite maturation compared to uninfected host cells. P1 and P2 spores population production were monitored by flow cytometry within the same culture every 30 minutes from 35.5 to 39 hours. P2 was the first spore type to emerge from infection, followed by P1 population after 1.5 hours. **I** = infected hosts, H = heath hosts; P1, P2 = spore populations.

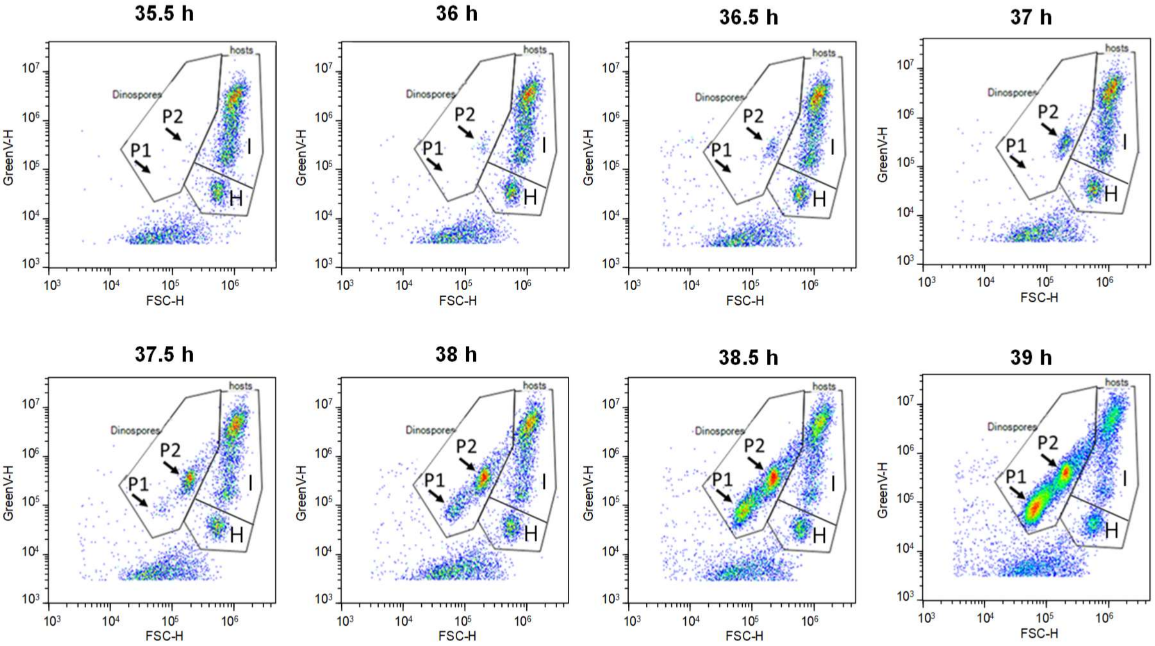
