## Supplementary Figure 2 for "Two Spore Types in a Marine Parasite of Dinoflagellates"

**Supplementary Figure 2**: Flow cytometry analysis of spore types released from individually picked infected host cell. The infected culture used for cell isolation released as much P1 as P2 over the infection cycle (A, 54 h post infection). Out of a total of 45 sorted cells, nearly all infected host cells produced exclusively either P1 (C, 6 isolated cells with a similar release) or P2 (E, 35 isolated cells) spores. A greenish P1-like population was also observed in two isolated cells (B). Two additional infected host produced either P1 spores and larger cells (D) and P2 spores and larger cells (F). Spores are produced from a multicellular sporont, called the vermiform stage, released directly from the infected host. This stage has a very short lifespan, typically ranging from a few minutes to a few hours. Each cell separates, divides, giving rise to one to several final spores. We hypothesized that the larger populations were composed of immature spores.

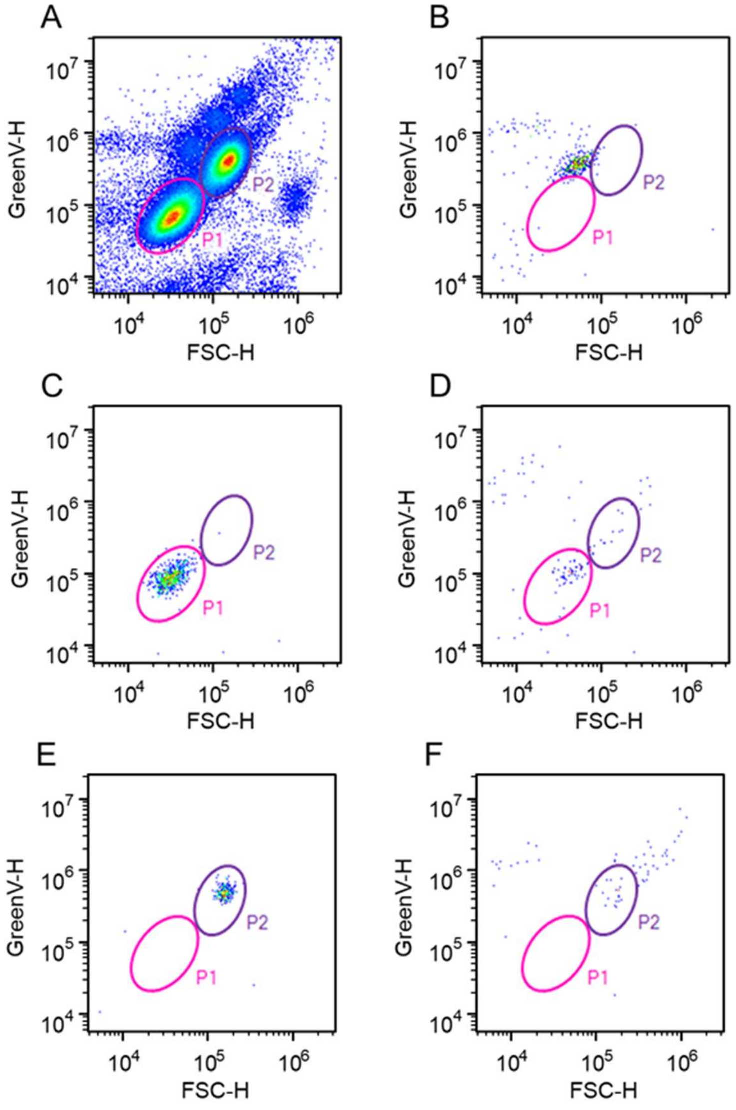
