## Supplementary Figure 3 for "Two Spore Types in a Marine Parasite of Dinoflagellates"

**Supplementary Figure 3:** Fate of distinct spore populations (P1, P2, and P3) tracked by flow cytometry, measured in cells per mL over time during two distinct experiments: A) populations followed from 25 to 45 hours post-inoculation, B) populations followed from 60 to 140 hours post-inoculation. P3 spores were produced 50 hours post-inoculation. The density of P1 remained relatively stable over time, while both P2 and P3 rapidly declined after 70 hours. P2 is produced slightly before P1.

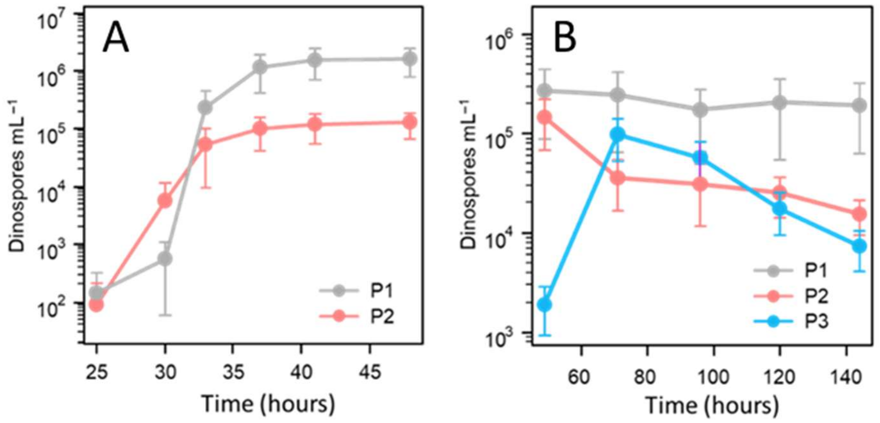
