## Supplementary Figure 4 for "Two Spore Types in a Marine Parasite of Dinoflagellates"

**Supplementary Figure 4: Spore biovolumes estimated by confocal microscopy.** (A) Acquisition from sample 1, dominated by P1 spores (FCM: 88%, Confocal microscopy: 86.6%); (B) Acquisition from sample 2, dominated by P2 spores (FCM: 85%, Confocal microscopy: 87%); (C) Bimodal distribution of cell sizes from sample 1 and 2, assessed by volume measurements from 3D image confocal microscopy.

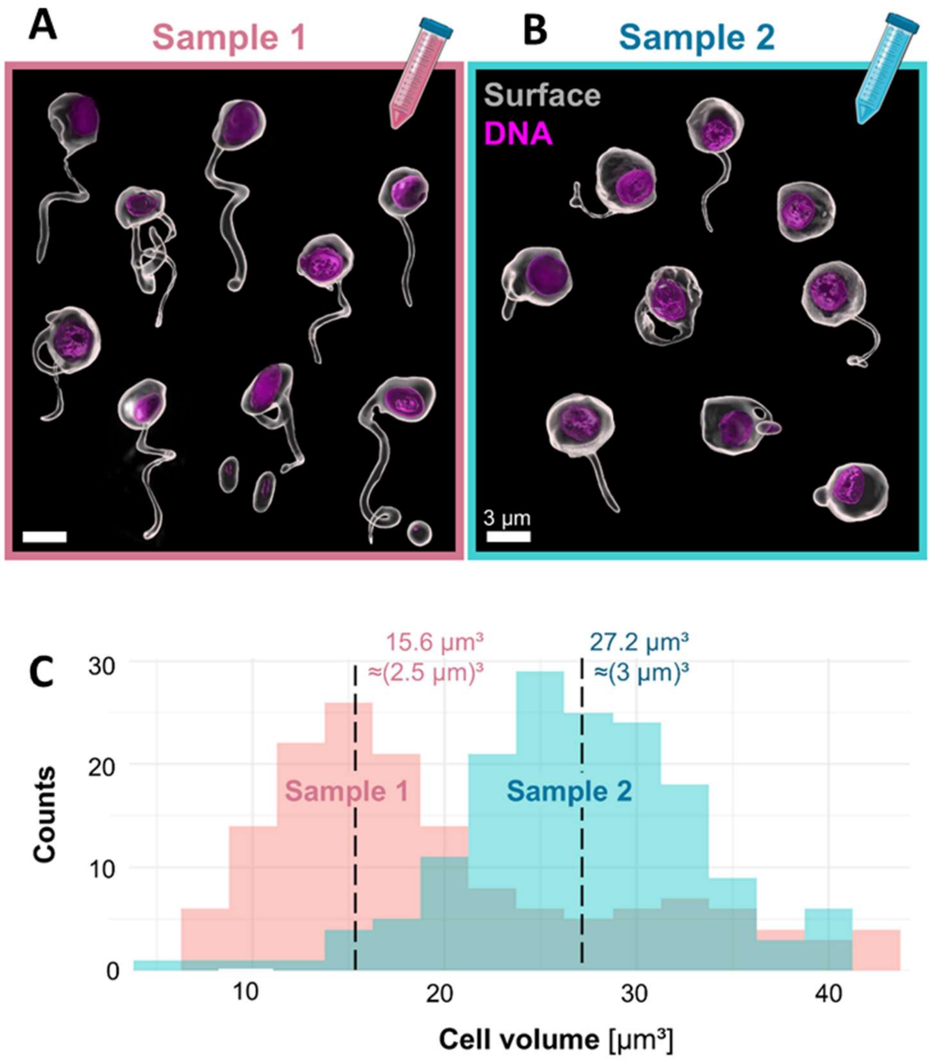
