## Supplementary Figure 5 for "Two Spore Types in a Marine Parasite of Dinoflagellates"

**Supplementary Figure 5: Infectivity of P2 spores monitored by flow cytometry.** Healthy and infected host are on the right, spores on the left (P1, P2, and P3). No infection was observed in host (followed during a week), while P2 population mainly transformed into P3 at t=45 hours post-inoculation.

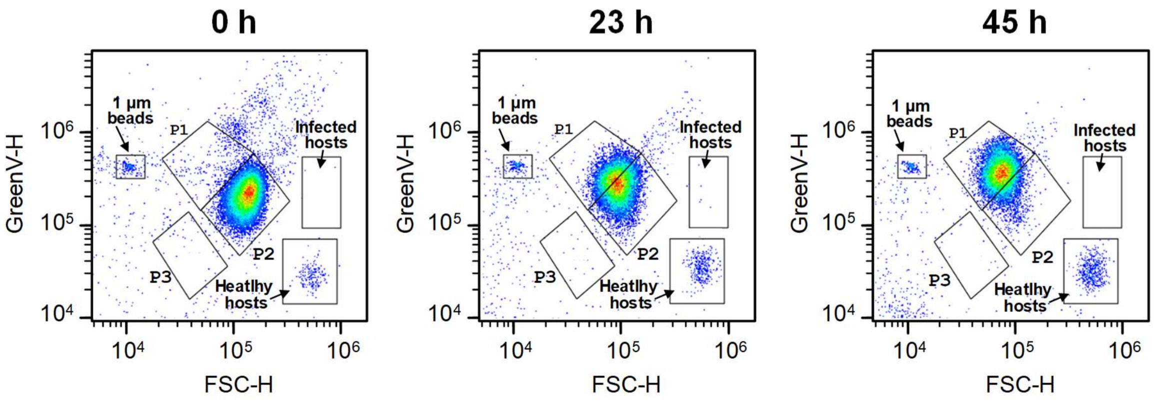
