## Supplementary Figure 6 for "Two Spore Types in a Marine Parasite of Dinoflagellates"

**Supplementary Figure 6: Ploidy level of mixed populations of spores as observed by flow cytometry. The left panel** (A, D) shows a density plot of living spore cultures, illustrating their natural distinct autofluorescence (Green-V, 405 nm laser) and size characteristics (FSC-H). Moving to the **middle panel** (B, E), we observe a contour plot of extracted nuclei after NIB/2 treatment stained by SYBR green I. In this plot, the nuclei are differentiated based on the green fluorescence intensity after SYBR green I staining (Green-B-H, 488 nm laser) and size (FSC-H). The **right panel** (C, F) displays the same nuclei as shown in the middle panel, visualized on a histogram. This histogram is based on density (counts) and green fluorescence (Green-B-H). **The upper layer** (A-C) represents a spore culture consisting of 58% P1 and 40% P2. The ratio of the P1 and P2 nuclei fluorescence is 1.17. **The lower layer** (D-F) represents a spore culture comprising 76% P1 and 20% P3. The ratio of the P1 and P3 nuclei fluorescence is 1.18. This ratio should be close to 2 in the case of a doubling of the ploidy level, which is not the case. We concluded that the ploidy level is similar, with the slight difference possibly due to the difference in chromatin conformation.

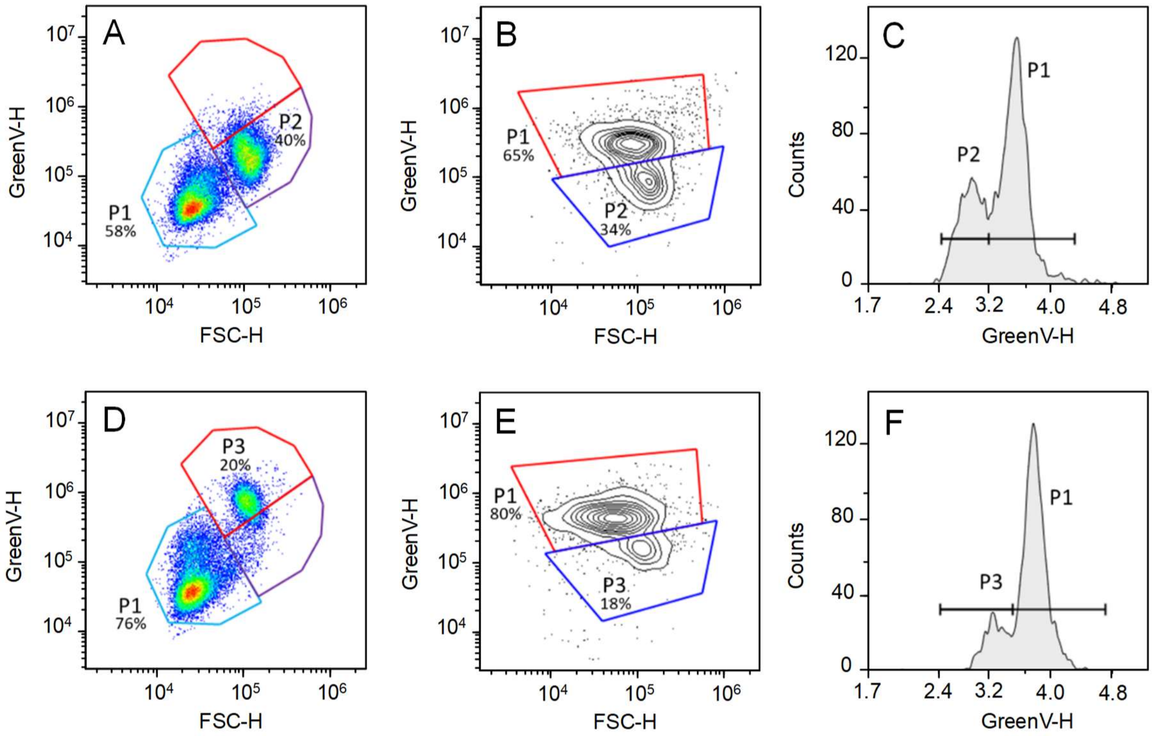
