## Supplementary Figure 7 for "Two Spore Types in a Marine Parasite of Dinoflagellates"

**Supplementary Figure 7: Populations sorted by flow cytometry for the analysis of their gene expression (transcriptomic analyses).** (A) Gates used for sorting infected host cell (20 cells per sample), and (B-C) gates used for sorting spores (10 cells per sample) for transcriptomic analyses. Different spore populations were gated using the 405 nm laser (green autofluorescence) and FSC. For host cell populations (uninfected and infected host cells), two successive gating strategies were used. The whole microalgal population was initially gated using the red autofluorescence under 488 nm excitation against FSC. Subsequently, a second gating strategy allowed for sorting host cells at different stages of the infection cycle, from the green fluorescence signal of the parasite measured under 405 nm excitation.

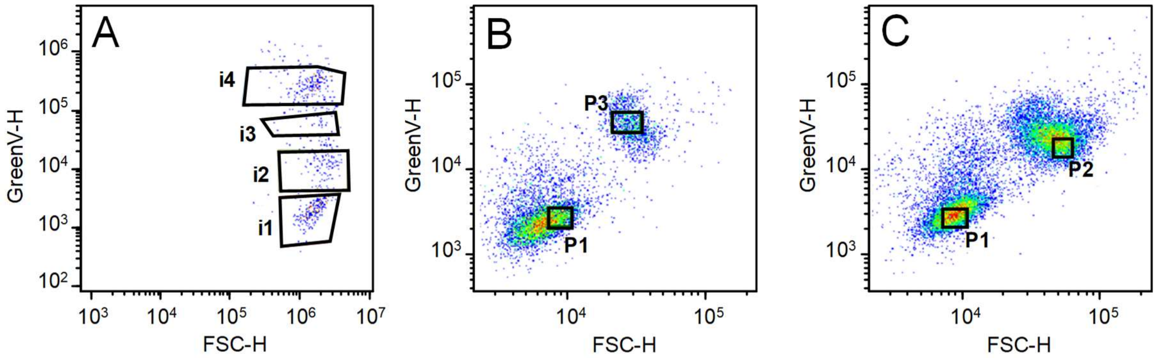
