## Supplementary Figure 8 for "Two Spore Types in a Marine Parasite of Dinoflagellates"

**Supplementary Figure 8: Heatmap analyses of gene expression along the course of infection (four stages of infection, namely i1, i2+i3, i4) and in spore populations (P1 or P2+P3) for selected metabolic pathways (genes involved in the earlier development of ribosomes, DNA and RNA synthesis, and cell replication).**

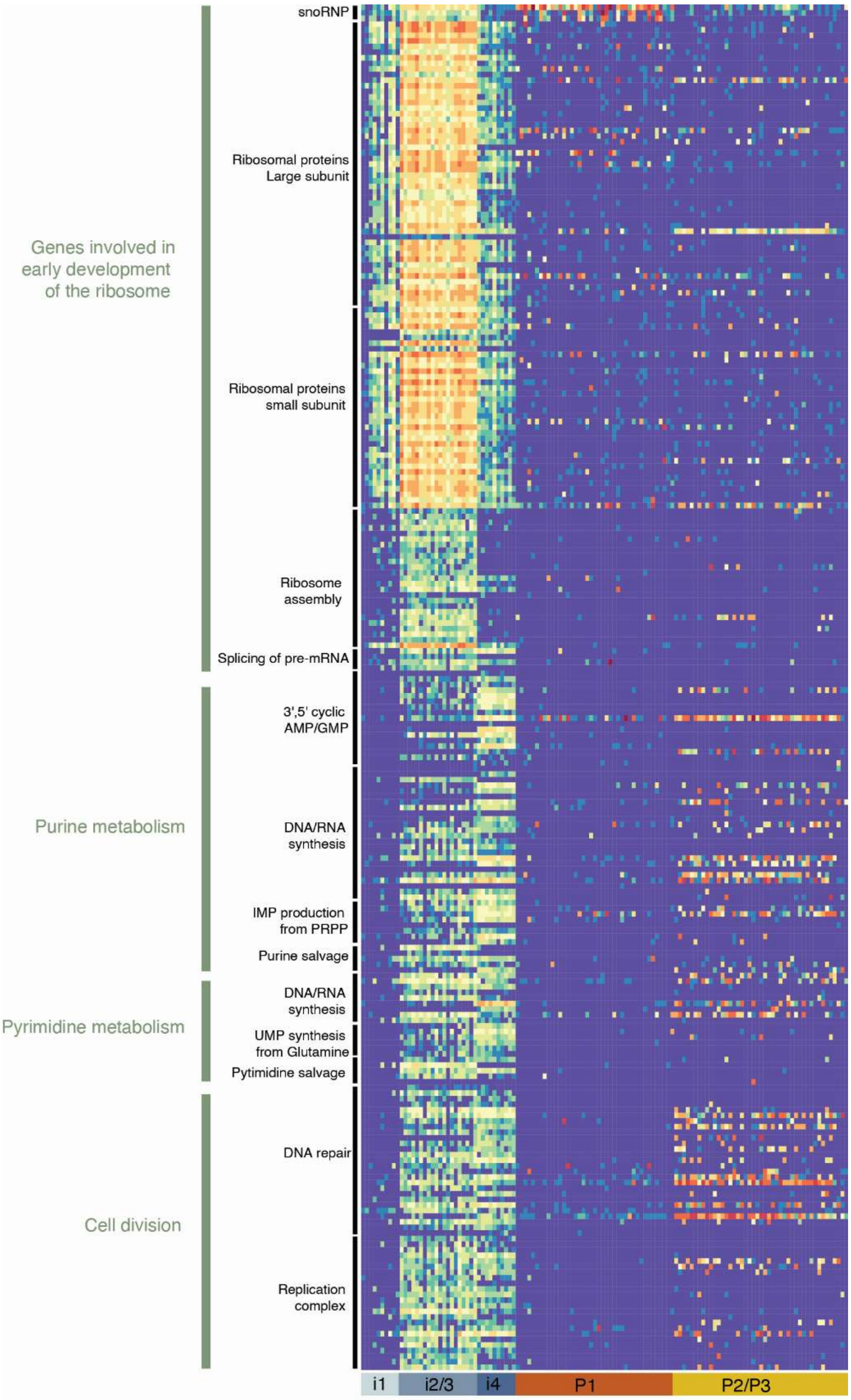
