## Supplementary Figure 9 for "Two Spore Types in a Marine Parasite of Dinoflagellates"

**Supplementary Figure 9**: **Exometabolomic analyses.** A- Principal component analysis was conducted on the 272 selected features specifically associated with P1 or P2 spore cell exudate extract profiles. B- Top 50 significant compounds discriminating the exometabolome profiles of P1 and P2 extracts were annotated. The intensities detected for the 50 metabolites were square root-transformed and Pareto-scaled, then displayed in the heatmap. 1-day old corresponds to t=49 h post-inoculation, 2-day old corresponds to t=71 h post-inoculation.

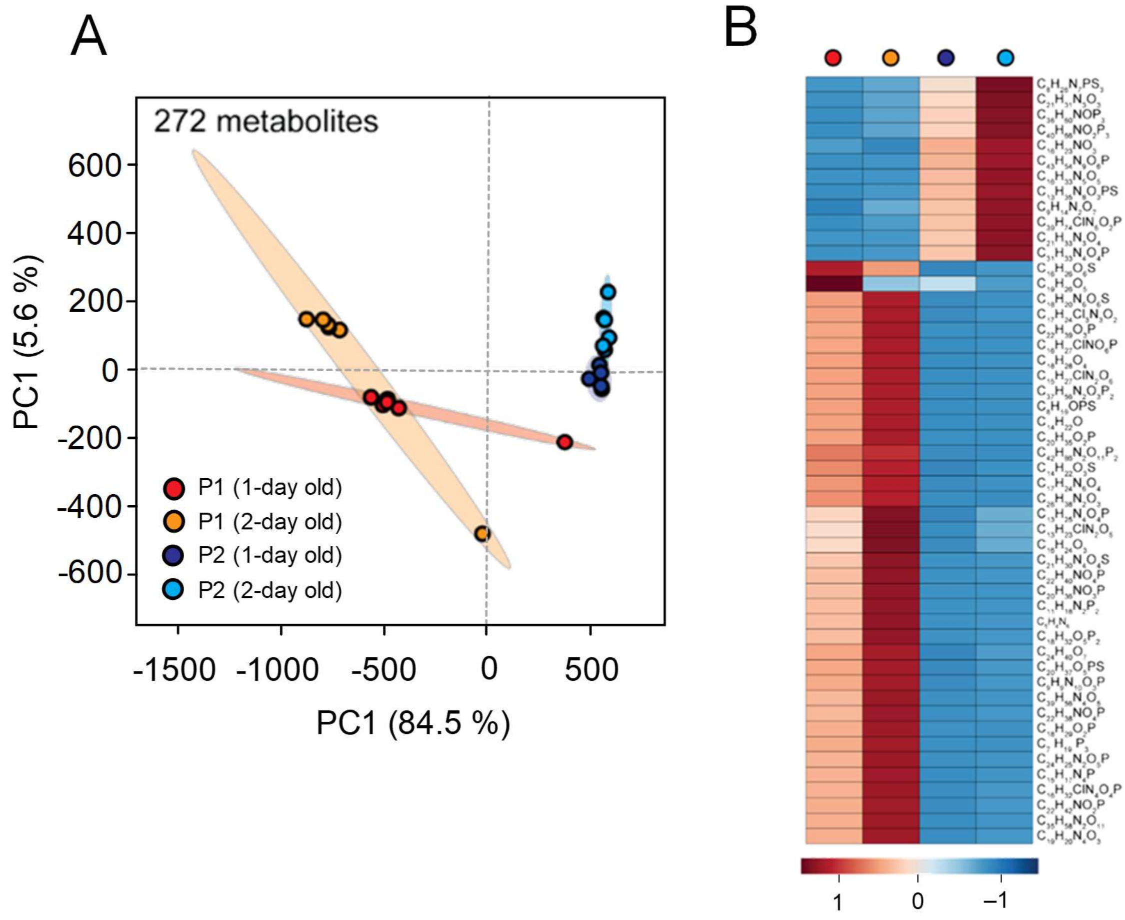


**Extended Methods supporting Supplementary Figure 9**

*Metabolic extractions, UHPLC-HRMS profiling, data-dependent MS acquisition and metabolites annotation*

We followed the protocols outlined in Vallet et al., (2019) for cell and exudate extraction and analysis using comparative metabolomics. Two days after inoculation, 40 mL of spore cultures were vacuum-filtered using 25 mm GF/C microfiber filters (Whatman plc, Maidstone, UK). The GF/C filters were transferred directly to 2 mL safe-lock Eppendorf tubes and extracted with 1.6 mL cold (–20°c) methanol (99.8 %, anhydrous, SIGMA-ALDRICH Chemie GmbH, Munich, Germany) through sonication for 15 min in an ultrasonic cleaner Emmi-D280 (EMAG-AG, Herford, Germany). Particles were removed by pipetting the supernatant after two successive centrifugations (10 min at 10,000× g). The organic phases were transferred to new glass vials and dried using a vacuum concentrator (Speedvac, Thermo Ficher Scientific). The dried samples were stored at –20 °C until endometabolome profiling. The particle-free filtrate was extracted using the solid-phase extraction method (6cc OASIS HLB sorbent from Waters) with the following sequential steps: conditioning (3 mL of methanol HPLC grade), equilibration (3 mL of pure milliQwater), sample extraction (about 35 mL), salt washing (4 mL of pure milliQwater), and elution of the extract (3 mL of methanol). The eluted solvent was then evaporated at a temperature of 35°C using a vacuum concentrator (Speedvac, Thermo Ficher scientific) in glass tubes to obtain dry extracts for exometabolome profiling. For the UHPLC-HRMS analysis, the dried samples were suspended in 100 µL of methanol/water (1/1) and centrifuged for 20 minutes at 14,000 × g. Subsequently, 50 µL of each sample was transferred to 1.5 mL glass vials with inserts, and 10 µL per sample was pooled into a QC mix sample, excluding blanks. Additionally, 1 µL of the internal standard L-fluorophenylalanine (55 µM in milliQwater) was added to each sample, thus reaching a final concentration of 0.055 mM. QC blank samples were prepared by combining 10 µL of each blank sample from extracts of the axenic medium into one vial. 10 µL of all samples were injected into the UHPLC-HR-MS system, consisting of a Dionex UltiMate 3000® (Dionex, Germering, Germany) coupled with a Q-Exactive Plus Orbitrap mass spectrometer (Thermo Scientific, Bremen, Germany). Metabolite separation was achieved using a 12-minute gradient on an Accucore C18 column (100 × 2.1 mm, particle size of 2.6 µm, Thermo Scientific Fisher, Dreieich, Germany), starting with 100% aqueous phase (2% acetonitrile, 0.1% formic acid in water) and increasing the acetonitrile phase over 8 minutes until reaching 100%. This ratio was held for 3 min before switching back to 100% aqueous phase and equilibration for 1 min. The flow rate was set at 0.4 mL $min^{-1}$, and the column oven temperature was maintained at 25 °C. Mass spectrometry analysis was performed in positive and negative modes with a scan range of *m*/*z* 75 to 1125 and a peak resolution of 70,000 for MS1 acquisition. Electrospray ionization was carried out with the following parameters: capillary temperature of 380 °C, spray voltage of 3000 V, sheath gas flow of 60 arbitrary units, and auxiliary gas flow of 20 arbitrary units. For MS2 acquisition using ddMS TopN experiments, MS2 spectra were obtained at a peak resolution of 70,000 (NCE 15, 30, 45), using an AGC target set to $3 \times 10^6$ and a maximum ion time set to 100 ms. The MS/MS spectra of precursor ions were obtained from the pooled QC sample using the abovementioned MS parameters and with an isolation window of *m/z* 0.4.

LC-MS runs were visualized using the Xcalibur software (Thermo Fisher Scientific). Metabolome analysis, data processing, and peak deconvolution were performed using Compound Discoverer versionTM software (v 3.3.2.31; Thermo Fisher Scientific, Bremen, Germany) following a comparative untargeted metabolomics workflow. The raw data were imported for peak deconvolution and metabolite annotation. The mass tolerance for MS identification was set at 5 ppm, the minimum MS peak intensity was $2 \times 10^5$, and the intensity tolerance for the isotope search was 30 %. The relative standard deviation was set to 50%. The selected labeled spectra were exported as .xlsx files, and the masses were searched in public mass lists (LipidsMaps, Natural Products Atlas, Thermo libraries). The raw dataset was uploaded on MetaboLights [MTBLS6476] (https://www.ebi.ac.uk/metabolights/editor/MTBLS6476/descriptors). The compound list was exported as a .csv file, and the intensities were normalized based on a normalization factor determined by the total carbon content. PCA was performed to compare the similarities of metabolites between cellular extracts of *Amoebophrya* spores using MetaboAnalyst 5.0. (URL: https://www.metaboanalyst.ca/ accessed on 12 December 2022). The Interquartile filter was applied for all processed datasets, and the intensities were log-transformed and Pareto-scaled. The identity of selected compounds was further confirmed using tandem mass spectrometry, and the MS/MS spectra were compared by spectral analysis and similarity search using CSI:FingerID in SIRIUS (Dührkop et al., 2015).

Dührkop K, Fleischauer M, Ludwig M, Aksenov AA, Melnik AV, Meusel M, Dorrestein PC, Rousu J, Böcker S. SIRIUS 4: a rapid tool for turning tandem mass spectra into metabolite structure information. Nat Methods. 2019 Apr;16(4):299-302. doi: 10.1038/s41592-019-0344-8. Epub 2019 Mar 18. PMID: 30886413.

Vallet, M.; Baumeister, T.U.H.; Kaftan, F.; Grabe, V.; Buaya, A.; Thines, M.; Svatoš, A.; Pohnert, G. The oomycete *Lagenisma coscinodisci* hijacks host alkaloid synthesis during infection of a marine diatom. Nat. Commun. **2019**, 10, 4938.
