## Supplementary Figure 10 for "Two Spore Types in a Marine Parasite of Dinoflagellates"

**Supplementary Figure 10: Endometabolomic analyses.** A- Principal component analysis was conducted on the 44 selected metabolites specifically associated with spore cell exudate extract profiles at different age. B- The top 44 significant compounds discriminating the endometabolome profiles of P1 and P2 extracts were annotated. The intensities detected for the 44 metabolites were square root-transformed and Pareto-scaled, then displayed in the heatmap. * Molecules also detected in the exometabolome (putatively secreted molecules). Only molecules that were unambiguously identified with a standard are named. 1-day old corresponds to t=49 h, 2-day old corresponds to t=71 h.

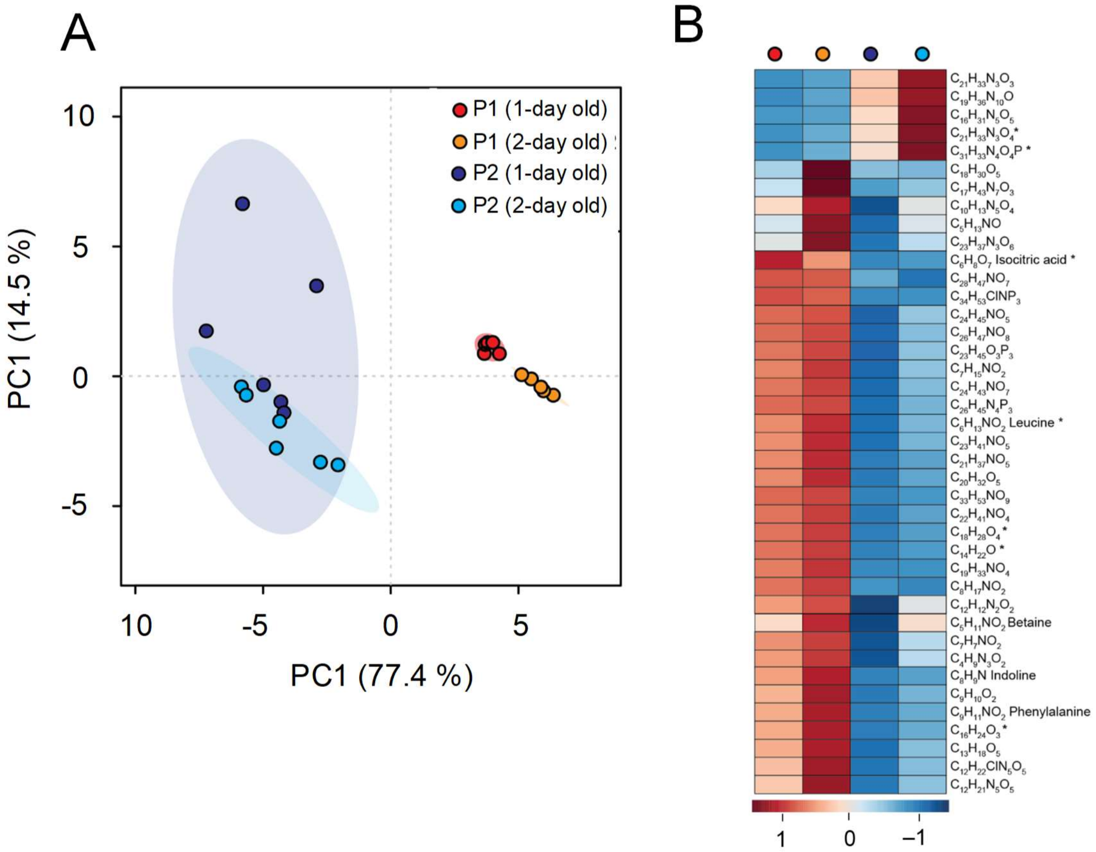


**Extended Methods supporting Supplementary Figure 10**

*Metabolic extractions, UHPLC-HRMS profiling, data-dependent MS acquisition and metabolites annotation*

We followed the protocols outlined in Vallet et al., (2019) for cell and exudate extraction and analysis using comparative metabolomics. Two days after inoculation, 40 mL of spore cultures were vacuum-filtered using 25 mm GF/C microfiber filters (Whatman plc, Maidstone, UK). The GF/C filters were transferred directly to 2 mL safe-lock Eppendorf tubes and extracted with 1.6 mL cold (–20°c) methanol (99.8 %, anhydrous, SIGMA-ALDRICH Chemie GmbH, Munich, Germany) through sonication for 15 min in an ultrasonic cleaner Emmi-D280 (EMAG-AG, Herford, Germany). Particles were removed by pipetting the supernatant after two successive centrifugations (10 min at 10,000× g). The organic phases were transferred to new glass vials and dried using a vacuum concentrator (Speedvac, Thermo Ficher Scientific). The dried samples were stored at –20 °C until endometabolome profiling. The particle-free filtrate was extracted using the solid-phase extraction method (6cc OASIS HLB sorbent from Waters) with the following sequential steps: conditioning (3 mL of methanol HPLC grade), equilibration (3 mL of pure milliQwater), sample extraction (about 35 mL), salt washing (4 mL of pure milliQwater), and elution of the extract (3 mL of methanol). The eluted solvent was then evaporated at a temperature of 35°C using a vacuum concentrator (Speedvac, Thermo Ficher scientific) in glass tubes to obtain dry extracts for exometabolome profiling. For the UHPLC-HRMS analysis, the dried samples were suspended in 100 µL of methanol/water (1/1) and centrifuged for 20 minutes at 14,000 × g. Subsequently, 50 µL of each sample was transferred to 1.5 mL glass vials with inserts, and 10 µL per sample was pooled into a QC mix sample, excluding blanks. Additionally, 1 µL of the internal standard L-fluorophenylalanine (55 µM in milliQwater) was added to each sample, thus reaching a final concentration of 0.055 mM. QC blank samples were prepared by combining 10 µL of each blank sample from extracts of the axenic medium into one vial. 10 µL of all samples were injected into the UHPLC-HR-MS system, consisting of a Dionex UltiMate 3000® (Dionex, Germering, Germany) coupled with a Q-Exactive Plus Orbitrap mass spectrometer (Thermo Scientific, Bremen, Germany). Metabolite separation was achieved using a 12-minute gradient on an Accucore C18 column (100 × 2.1 mm, particle size of 2.6 µm, Thermo Scientific Fisher, Dreieich, Germany), starting with 100% aqueous phase (2% acetonitrile, 0.1% formic acid in water) and increasing the acetonitrile phase over 8 minutes until reaching 100%. This ratio was held for 3 min before switching back to 100% aqueous phase and equilibration for 1 min. The flow rate was set at 0.4 mL $min^{-1}$, and the column oven temperature was maintained at 25 °C. Mass spectrometry analysis was performed in positive and negative modes with a scan range of *m*/*z* 75 to 1125 and a peak resolution of 70,000 for MS1 acquisition. Electrospray ionization was carried out with the following parameters: capillary temperature of 380 °C, spray voltage of 3000 V, sheath gas flow of 60 arbitrary units, and auxiliary gas flow of 20 arbitrary units. For MS2 acquisition using ddMS TopN experiments, MS2 spectra were obtained at a peak resolution of 70,000 (NCE 15, 30, 45), using an AGC target set to $3 \times 10^6$ and a maximum ion time set to 100 ms. The MS/MS spectra of precursor ions were obtained from the pooled QC sample using the abovementioned MS parameters and with an isolation window of *m/z* 0.4.
