## Supplementary Table 1 for "Two Spore Types in a Marine Parasite of Dinoflagellates"

**Supplementary Table 1: Gene orthologues in *Amoebophrya* genome strain A120 for genes involved in meiosis**. *meiosis-specific genes. ND=not detected.

Reference sequences were used as BLAST queries to identify homologs in the Amoebophrya genome (available here: http://application.sb-roscoff.fr/blast/hapar/download.html), and the identity of positive hits was confirmed by (1) reverse-BLAST to the UniProtKB database (https://www.uniprot.org/blast/; last access September 2023); (2) sequence search in InterPro (http://www.ebi.ac.uk/interpro/; last access September 2023); (3) domain search with Pfam 34.0 (http://pfam.xfam.org/; last access September 2023); (4) phylogeny performed on the NGPhylogeny.fr website (https://ngphylogeny.fr/)[1] where genes were aligned using mafff v.7[2] alignments, filtered using trimAl[3], and Maximum Likelihood (ML) trees were constructed using FastTree v.2[4,5] with 1000 bootstraps for branch support. The homology of genes was based on visual inspection of alignments using SeaView v.5.0.5[6] and of phylogenetic trees using FigTree v1.4.4 (http://tree.bio.ed.ac.uk/software/figtree/). The final set of homologous sequences were then aligned with mafft online[7], and the alignments were filtered with Gblocks v. 0.91b with the -b5=a option[8]. Single gene phylogenetic trees were reconstructed for each alignment using RAxML v. 8.2.12[9] with the -# 1000 -m PROTGAMMAIAUTO options. The latter phylogenetic analyses were performed on the ABiMS platform (http://abims.sb-roscoff.fr/) at the Station Biologique de Roscoff.


1. Lemoine, F. et al. NGPhylogeny.fr: new generation phylogenetic services for non-specialists. Nucleic Acids Research 47, W260–W265 (2019).
2. Katoh, K. & Standley, D. M. MAFFT Multiple Sequence Alignment Software Version 7: Improvements in Performance and Usability. Molecular Biology and Evolution 30, 772–780 (2013).
3. Capella-Gutiérrez, S., Silla-Martínez, J. M. & Gabaldón, T. trimAl: a tool for automated alignment trimming in large-scale phylogenetic analyses. Bioinformatics 25, 1972–1973 (2009).
4. Price, M. N., Dehal, P. S. & Arkin, A. P. FastTree: Computing Large Minimum Evolution Trees with Profiles instead of a Distance Matrix. Molecular Biology and Evolution 26, 1641–1650 (2009).
5. Price, M. N., Dehal, P. S. & Arkin, A. P. FastTree 2 – Approximately Maximum-Likelihood Trees for Large Alignments. PLoS ONE 5, e9490 (2010).
6. Gouy, M., Tannier, E., Comte, N. & Parsons, D. P. Seaview Version 5: A Multiplatform Software for Multiple Sequence Alignment, Molecular Phylogenetic Analyses, and Tree Reconciliation. in Multiple Sequence Alignment (ed. Katoh, K.) vol. 2231 241–260 (Springer US, New York, NY, 2021).

7. Katoh, K., Rozewicki, J. & Yamada, K. D. MAFFT online service: multiple sequence alignment, interactive sequence choice and visualization. Brief. Bioinformatics 20, 1160–1166 (2019).
8. Castresana, J. Selection of conserved blocks from multiple alignments for their use in phylogenetic analysis. Mol. Biol. Evol. 17, 540–552 (2000).
9. Stamatakis, A. RAxML version 8: a tool for phylogenetic analysis and post-analysis of large phylogenies. Bioinformatics 30, 1312–1313 (2014).

| Processes | Name | Symbol | A120_Gene_ID | REF |
|---|---|---|---|---|
| Induction | switch from mitotic to m | Meio, Mei2 (in vegetative cells after3) | GSA120T00008208001 | This study |
| Mismatch corre | DNA mismatch repair | Msh2 (in vegetative cells after3) | GSA120T00021256001 | Cai et al. 2019 |
| | DNA mismatch repair | Msh3 | ND | Cai et al. 2019 |
| | DNA mismatch repair | Msh6 (in vegetative cells after3) | GSA120T00017079001 | Cai et al. 2019 |
| | DNA mismatch repair (m | PMS1 | ND | This study |
| | DNA mismatch repair (m | PMS2 (in resting cysts after3) | GSA120T00014436001 | This study |
| Syngamy | Plasmogamy | Hap2* (in vegetative cells after3) | GSA120T00024727001 | This study |
| | | **GEX1*** | ND in dinoflagellates | Shah et al. 2020 |
| Cohesin complex | | REC8 | ND in dinoflagellates | Shah et al. 2020, Cai et |
| | Cohesion complex | Smc1 (in resting cysts after3) | GSA120T00000722001 | This study |
| | | Smc1 (in resting cysts after3) | GSA120T00021886001 | This study |
| | | Smc2 | GSA120T00011437001 | This study |
| | | Smc4 | GSA120T00003312001 | This study |
| | | Smc4 | GSA120T00003311001 | This study |
| | | Smc3 | GSA120T00016095001 | This study |
| | | Smc5 | ND in dinoflagellates | Shah et al. 2020 |
| | | Smc6 | ND in dinoflagellates | Shah et al. 2020 |
| SC formation | | **Hop1*** | ND in dinoflagellates | Shah et al. 2020 |
| | | **Red1*** | ND in dinoflagellates | Shah et al. 2020 |
| | | **Pch2*** | ND in dinoflagellates | Shah et al. 2020 |
| ZMM protein | | **Msh4* (in gametes after3)** | ND | Cai et al. 2019 |
| | | **Msh5* (in resting cysts after3)** | ND | Cai et al. 2019 |
| | | **Mer3*** | ND | Cai et al. 2019 |
| | | Zip1 | ND in dinoflagellates | Shah et al. 2020 |
| | | Zip2 | ND in dinoflagellates | Shah et al. 2020 |
| | | Zip3 | ND in dinoflagellates | Shah et al. 2020 |
| Double strande | Meiotic recombination p | **SPO11* (in germinating cysts after3)** | GSA120T00013268001 | Cai et al. 2019 |
| | Meiotic recombination p | Mre11 (in resting cysts after3) | GSA120T00000265001 | This study |
| | | Dna2 | GSA120T00007919001 | This study |
| | | Atm | GSA120T00024825001 | This study |
| | Double strand breaks | RAD50 (in germinating cysts after3) | GSA120T00021321001 | This study |
| Homologous re | Stimulation of DMC1 act | MND1* (in germinating cysts after3) | GSA120T00008837001 | Cai et al. 2019 |
| | Recombinase with specif | **HOP2* (in germinating cysts after3)** | GSA120T00000161001 | Cai et al. 2019 |
| | DNA strand invasion | **RAD51* (in resting cysts after3)** | GSA120T00009824001 | Cai et al. 2019 |
| | Stimulation of RAD51 ac | RAD54 (in germinating cysts after3) | GSA120T00006516001 | This study |
| | Meiotic recombination p | **DMC1* (in germinating cysts after3)** | GSA120T00010080001 | Cai et al. 2019 |
| Cross over I | DNA mismatch repair | MLH1 (in vegetative cells after3) | GSA120T00014170001 | This study |
| | | MLH3 | ND | This study |
| | | SLX1 | GSA120T00019260001 | This study |
| | | SLX4 | ND | This study |
| | | SGS1 | GSA120T00024235001 | This study |
| | | EXO1 | GSA120T00008768001 | This study |
| | | EXO1 | GSA120T00005899001 | This study |
| Cross over II | | MUS81 | GSA120T00001489001 | This study |
| | | MMS4/EME1 | ND | This study |
| | Meiotically upregulated g | Mug157 (in resting cysts after3) | ND | This study |
| | | Plant-Msh1 | GSA120T00012118001 | This study |
| | Endonuclease | MustS2? (in germinating cysts after3) | ND | This study |
| | | Pms1 (in resting cysts after3) | GSA120T00014436001 | This study |
| | Cohesion complex | RAD21 | GSA120T00017233001 | Cai et al. 2019 |
| | | RAD21 | GSA120T00009901001 | Cai et al. 2019 |
| | Control of meiotic divisio | Mns1 | GSA120T00011473001 | This study |
